# Optimizing histatin 5: Effects of K13 and K17 substitutions on proteolytic stability and antifungal activity

**DOI:** 10.64898/2026.01.31.703050

**Authors:** Wright K. Makambi, Victoria L. Chiu, Lydia Kasper, Bernhard Hube, Amy J. Karlsson

**Affiliations:** Department of Chemical and Biomolecular Engineering, University of Maryland, College Park, USA; Department of Microbial Pathogenicity Mechanisms, Leibniz Institute for Natural Product Research and Infection Biology – Hans Knöll Institute, Jena, Germany; sInstitute of Microbiology, Friedrich-Schiller University, Jena, Germany

**Author notes:** Corresponding author:* Amy J. Karlsson, Department of Chemical and Biomolecular Engineering, University of Maryland, 4418 Stadium Drive, College Park, MD 20742.

**Keywords:** Antimicrobial peptide (AMP), antifungal peptide, aspartic protease, *Candida albicans*, histatin 5, peptides, protease, proteolytic stability, protein design, protein engineering

## Abstract

*Candida albicans* is an opportunistic human fungal pathogen found in the oral cavity. Human saliva contains a 24-amino acid peptide called histatin 5 (Hst5) that has activity against *C. albicans*, but degradation of Hst5 by secreted aspartyl proteases (Saps) produced by *C. albicans* and by salivary proteases can reduce its antifungal efficacy. Building on our previous work that identified K13 and K17 as important residues for stability and activity of Hst5, we systematically investigated amino acid modification at these sites. Modifications explored the influence of hydrophobicity, charge, polarity, size, and aromaticity on Hst5’s interaction with Saps and saliva. The K13R variant retained proteolytic stability and antifungal activity after incubation with Sap1, Sap2, Sap3, and Sap9, while other K13 variants generally had reduced stability and activity, emphasizing the importance of a positive charge at this position. At K17, substitutions generally enhanced proteolytic stability and improved antifungal activity after incubation with Saps. We introduced the normalized intact peptide (NIP) parameter as a tool for identifying Hst5 variants with improved stability in the presence of multiple Saps, and NIP revealed K17W as the most proteolytically stable variant overall. Additionally, we observed modest differences in peptide stability in saliva, and the K17W variant was the only variant that retained more activity than Hst5 following incubation with saliva. We further assessed the K17W variant’s ability to prevent biofilm formation and found it to be more effective than the parent peptide Hst5. Our findings highlight the interactions between the Hst5 K13 and K17 residues with Saps and saliva and provide a strong foundation for future Hst5 engineering efforts to improve proteolytic stability and antifungal efficacy in diverse proteolytic environments.

## Introduction

*Candida albicans* is a commensal fungal pathogen commonly found on mucosal surfaces, and it can cause an array of infections including oral thrush ^1-2^. Currently, small-molecule antifungal agents are used to treat *C. albicans* infections. However, due to the growing resistance of *C. albicans* and other pathogens to these agents ^3-4^, new antifungal agents are needed, and antifungal peptides are being explored as potential alternatives.

Histatin 5 (Hst5) is a 24 amino acid peptide secreted by the salivary glands, known for its activity against *C. albicans* including several drug-resistant strains ^5-8^. Unlike many antifungal peptides, which function by forming pores in cell membranes, Hst5 translocates into the fungal cell to exert its effects rather than simply disrupting the cell membrane. Several key events trigger this translocation: the accumulation of Hst5 on the surface of *C. albicans*, its interaction with the heat shock proteins Ssa1p and Ssa2p ^9^, and translocation via the polyamine transporters Dur3p and Dur31p ^10-12^. Due to its efficacy against *C. albicans*, Hst5 has been considered as a potential therapeutic.

While Hst5 shows promise as a therapeutic agent, its proteolytic stability remains a challenge. For example, *C. albicans* produces secreted aspartyl proteases (Saps) that can degrade Hst5. The Saps of *C. albicans* constitute a family of 10 enzymes, with Sap1-8 secreted fully to the extracellular space and Sap9 and Sap10 localized in the cell wall and cell membrane via glycosylphosphatidylinositol (GPI) anchors ^13-15^. Notably, prior studies comparing the protein expression of *C. albicans* in the planktonic and biofilm forms showed that *C. albicans* biofilms have increased expression of Saps, which creates a further challenge in using Hst5 as a therapeutic ^16-19^. Proteomic studies have revealed that Saps preferentially cleave near arginine, lysine, and hydrophobic residues ^14, 20^. Since the Hst5 sequence includes four lysine, three arginine, and three large hydrophobic amino acids, the peptide is prone to proteolysis by Saps.

Although we know what types of amino acids interact with Saps, little is known about the role different amino acid properties play in the degradation of Hst5 by Saps. In previous work from our laboratory, we identified important interactions between Hst5 and various Saps by substituting each lysine residue in Hst5 with either arginine or leucine ^21-22^. When incubated with recombinant Sap9, the K13L variant was more prone to proteolysis than the parent Hst5, while K17 variants (K17R and K17L) exhibited enhanced proteolytic stability. While these studies highlight the significance of the lysine residues in the Hst5-Sap interactions, only leucine and arginine modifications were explored, leaving the effect of other amino acid properties at K13 and K17 unknown.

If used as a treatment for oral candidiasis, Hst5 would be exposed not only to Saps but also to other proteases from host cells and microbial organisms present in the saliva ^23-25^. Prior studies demonstrated that Hst5 degrades upon exposure to saliva ^25-27^, and our earlier work showed that the Hst5 variant K17L has slightly improved proteolytic stability when incubated with saliva ^22^. The modest increase suggests that assessing only arginine and leucine substitutions may not be sufficient for identifying Hst5 variants that are stable in saliva, warranting further investigation of additional amino acid properties.

In this work, we explored how amino acid modifications at positions K13 and K17 of Hst5 affect antifungal activity and proteolytic stability in the presence of Sap1, Sap2, Sap3, Sap9, and saliva. We explored the amino acid properties that enhance or reduce the proteolytic stability and antifungal activity of Hst5 variants. Furthermore, we assessed the ability of the most stable peptide to prevent biofilm formation. Our results provide insight into peptide modifications that enhance the therapeutic potential of Hst5.

## Results

To investigate the effect of substitutions at K13 and K17 on proteolytic stability and the residual antifungal activity after degradation by Saps and saliva, we selected 12 Hst5 variants with substitutions at these sites (**Table 1**). We selected a range of substitutions to study the effect of hydrophobicity (K13A, K13L, K17A, K17L, and K17W), charge (K13E, K13H, K13R, K17E, and K17R), polarity (K13Q and K17Q), size (K13A, K17A, and K17W), and aromaticity (K13H and K17W). The breadth of the explored peptide library allowed us to study the effect of a range of amino acid side chain properties on Hst5’s proteolytic stability in the presence of fungal and salivary enzymes and antifungal activity. We tested the proteolytic stability in the presence of seven Saps: Sap 1, Sap2, Sap3, Sap5, Sap6, Sap9, and Sap10. Consistent with our previous results ^22^, Sap5, Sap6, and Sap10 did not degrade Hst5 or the variants (**Figure S1 and S2**), so our experiments focused on Sap1, Sap2, Sap3, and Sap9.

**Table 1.**
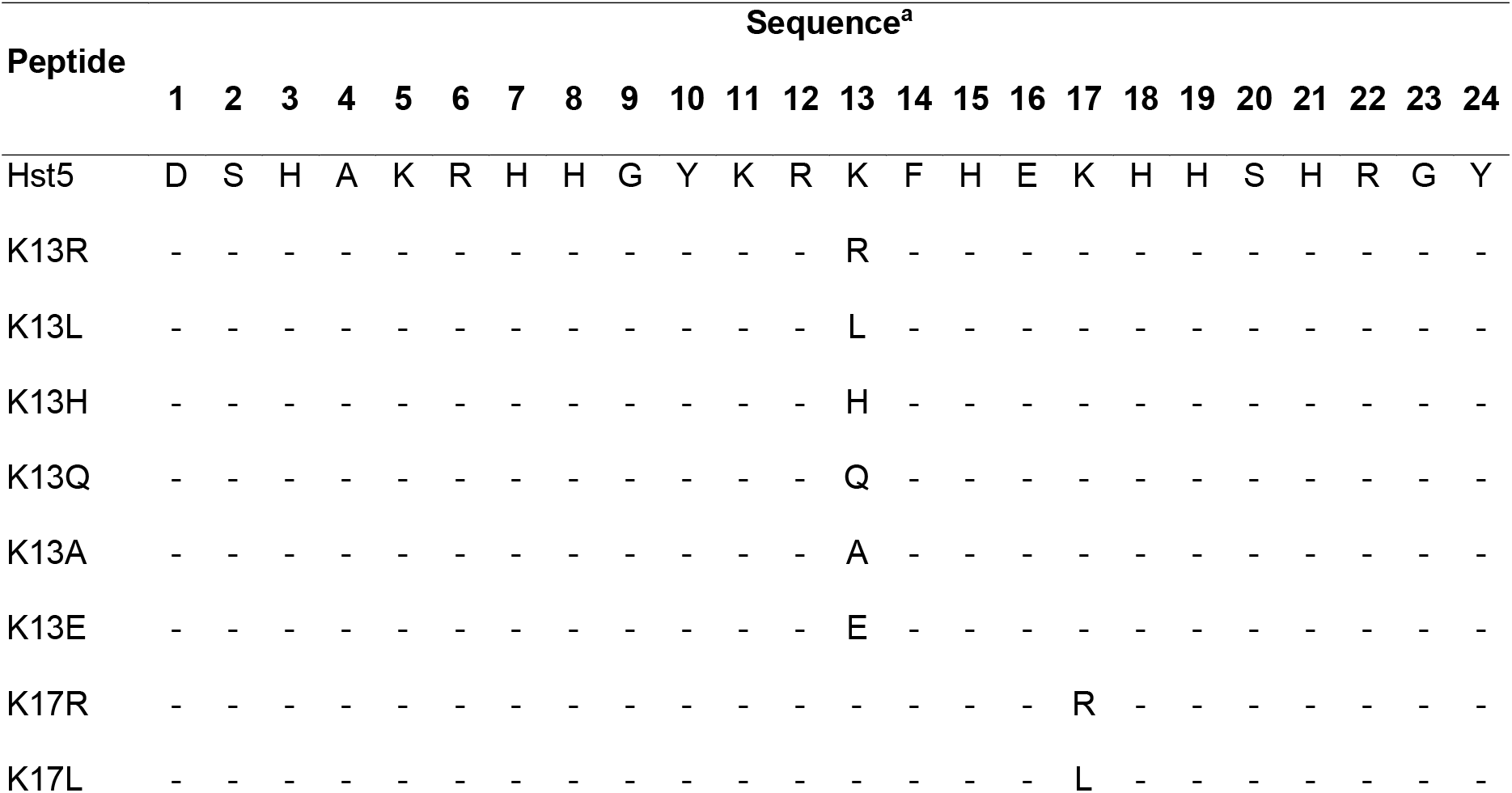
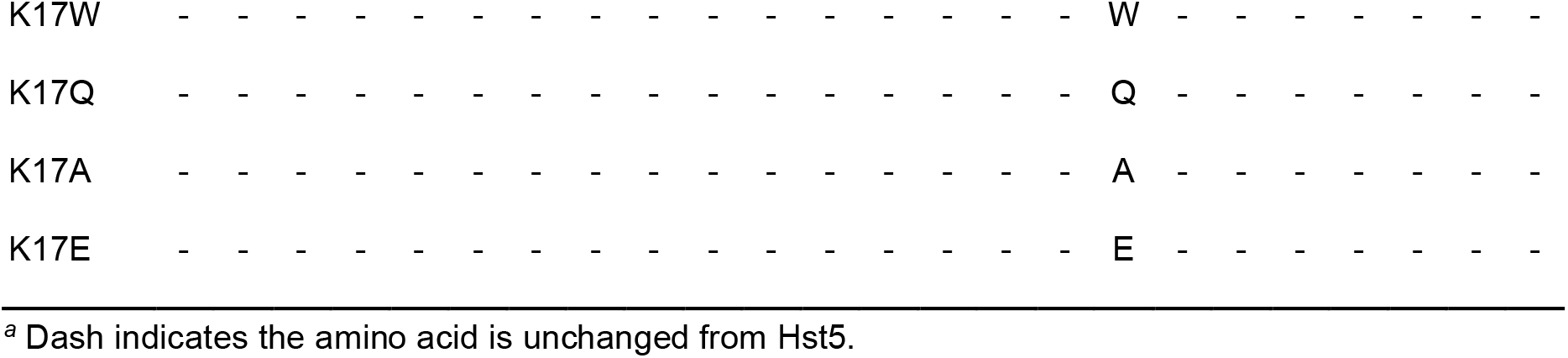
Peptide variants evaluated for proteolysis and antifungal activity.

### Proteolytic stability of Sap-treated K13 variants

To determine the extent of proteolysis for each peptide variant by the *C. albicans* Saps, we used gel electrophoresis and densitometry to quantify the intact peptide after incubation with recombinant Sap1, Sap2, Sap3, or Sap9. We selected concentrations of each Sap that preserved 25–50% of the fully intact Hst5 peptide after incubation with the Sap for two hours at 37 °C in 1 mM sodium phosphate buffer (NaPB) at pH 7.4. This strategy allowed for the clear identification of differences in the proteolytic stability among Hst5 and the peptide variants.

Under the conditions tested, Sap1, Sap2, Sap3, and Sap9 cleaved Hst5 and its variants with substitutions at K13, and the cleavage was dependent on the specific protease and the peptide modifications. After incubation with Sap1 (**Figure 1A**), 36.6% of Hst5 remained fully intact. Only K13R was as stable as the parent peptide in the presence of Sap1; all other variants were less stable. Incubation with Sap2 (**Figure 1B**) resulted in 49.2% of Hst5 remaining intact, and the only variant with enhanced stability in the presence of Sap2 was K13R, with 60.0% of the peptide remaining fully intact. All other variants had diminished stability, falling between 10.5%–26.8% of the peptide remaining intact. Incubation with Sap3 (**Figure 1C**) left 45.0% of Hst5 fully intact, and K13R was as stable as the parent peptide in the presence of Sap3. The K13E variants had enhanced proteolytic stability in the presence of Sap3, with 59.8% of the peptide remaining fully intact. The remaining variants were less stable, with K13H, K13Q, K13L, and K13A each having less than 5% of the intact peptide remaining. Finally, after incubation with Sap9 (**Figure 1D**), 41.6% of Hst5 remained fully intact, and K13R and K13A had comparable results. The remaining variants had reduced proteolytic stability, with between 5.2%–15.0% remaining intact. Our results indicate the importance of maintaining the positive charge at K13, as K13R was the only variant consistently as stable or more stable than the parent peptides Hst5.

**Figure 1.**
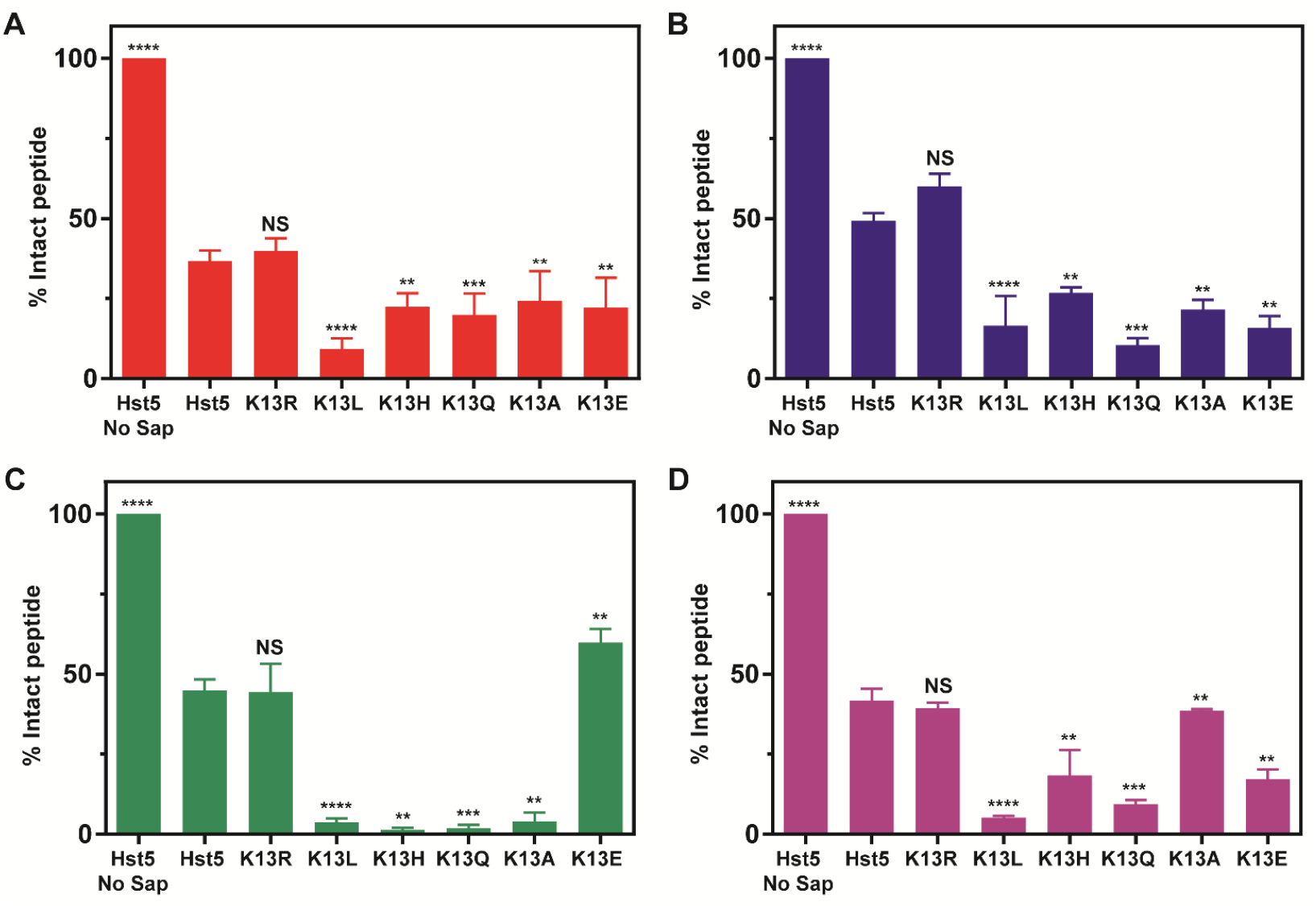
Degradation of Hst5 and K13 variants following incubation with purified recombinant A) Sap1, B) Sap2, C) Sap3, and D) Sap9. Peptides were incubated with the Saps for 2 h at 37 °C, and intact peptides and their degradation fragments were separated by gel electrophoresis. The percentage of peptide remaining intact was quantified by densitometry. Error bars represent the standard error of the mean (*n =* 6 for the Hst5 controls with and without Sap and *n =* 3 for the Hst5 variants). Representative gel images are provided in **Figure S3.**Statistical significance, relative to degraded Hst5, is indicated as follows: ns for not significant, * for p ≤ 0.05, ** for *p* ≤ 0.01, *** for *p* ≤ 0.005, and **** for *p* ≤ 0.0001.

### Proteolytic stability of Sap-treated K17 variants

While K13 modifications typically resulted in a reduction of proteolytic stability, modifications at K17 generally resulted in similar or improved proteolytic stability compared to the parent peptide after incubation with the Saps. After incubation with Sap1 (**Figure 2A**), the variants K17Q and K17E had enhanced proteolytic stability compared to Hst5, with 51.4% and 49.6% of the peptide remaining fully intact, respectively, while the stability of the remaining variants was similar to Hst5. Incubation with Sap2 produced more substantial differences compared to Hst5 (**Figure 2B)**. While the stability of K17E was statistically similar to Hst5 in the presence of Sap2, the K17R variant was less proteolytically stable, with only 35.1% of the peptide remaining intact. All other variants had enhanced proteolytic stability in the presence of Sap2, with 72.9%–81.1% of peptide remaining intact. For Sap3 (**Figure 2C**), the K17R and K17L modifications resulted in proteolytic stability similar to that seen for Hst5. All other K17 variants showed an improvement in proteolytic stability in the presence of Sap3, with 57.3%–66.9% of intact peptide remaining. Finally, after incubation with Sap9 (**Figure 2D**), all K17 variants had enhanced proteolytic stability with 68.1%– 79.0% of peptide remaining intact. Our findings indicate that substitutions away from K17 generally does not significantly change interactions with Sap1, substitution to uncharged residues improves stability in the presence of Sap2, and lysine at K17 can be replaced to enhance stability against Sap3 and Sap9.

**Figure 2.**
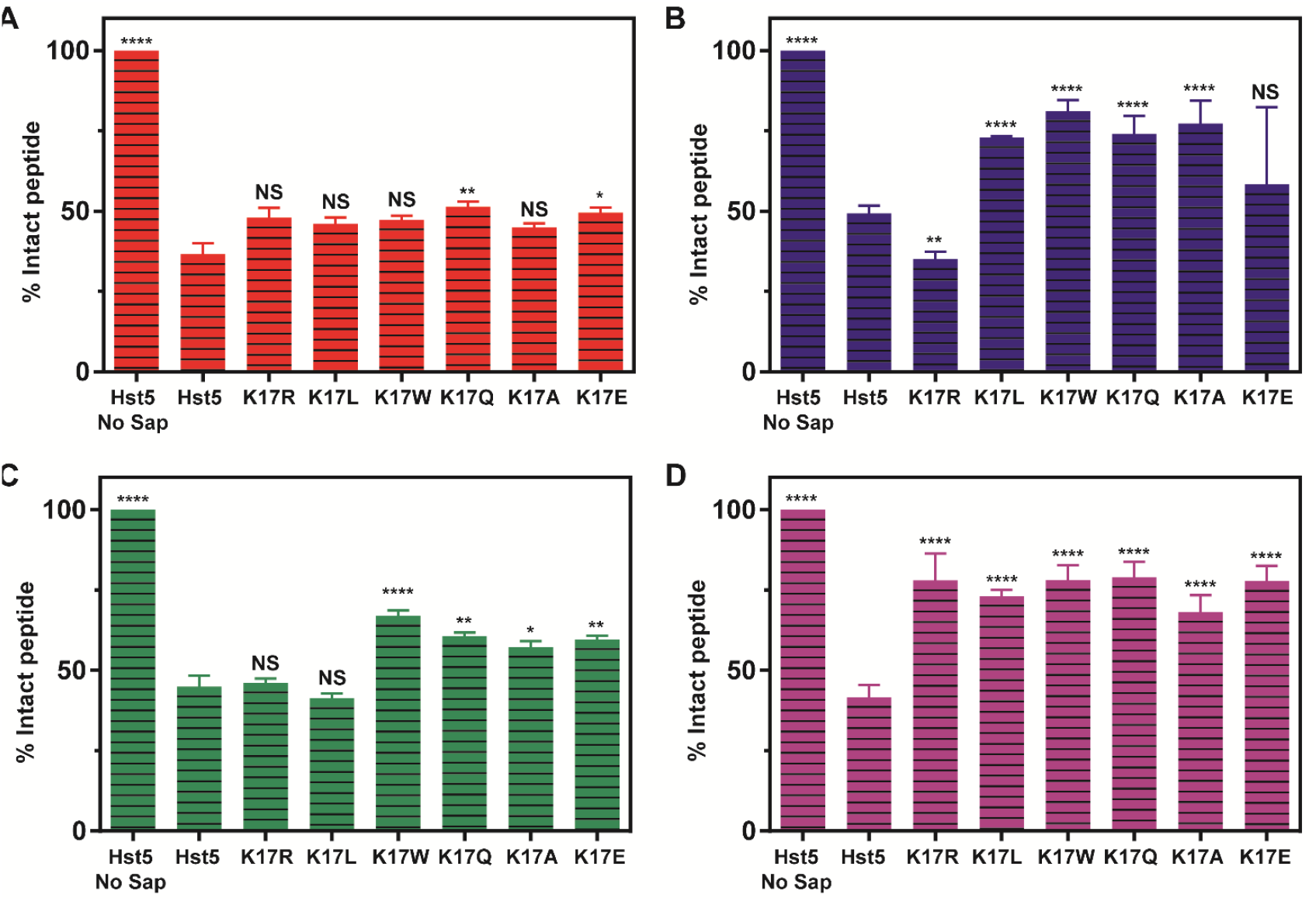
Degradation of Hst5 and K17 variants following incubation with purified recombinant A) Sap1, B) Sap2, C) Sap3, and D) Sap9. Peptides were incubated with the Saps for 2 h at 37 °C, and intact peptides and their degradation fragments were separated by gel electrophoresis. The percentage of peptide remaining intact was quantified by densitometry. Error bars represent the standard error of the mean (*n =* 6 for the Hst5 controls with and without Sap and *n =* 3 for the Hst5 variants). Representative gel images are provided in **Figure S4.**Statistical significance, relative to degraded Hst5, is indicated as follows: ns for not significant, ns for not significant, * for p ≤ 0.05, ** for *p* ≤ 0.01, *** for *p* ≤ 0.005, and **** for *p* ≤ 0.0001.

### Overall stability in the presence of Saps as determined by the normalized intact peptide

Because the peptides show varying levels of proteolytic stability across the different Saps, identifying the variant with the best overall stability compared to Hst5 is challenging. To address this, we developed the “normalized intact peptide (*NIP*)” parameter:

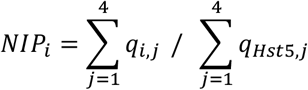

where *i* indicates the peptide variant, *j* indicates the specific Saps that degraded Hst5 and the variants (i.e., *j* = 1 for Sap1, *j* = 2 Sap2, *j* = 3 Sap3, or *j* = 4 for Sap9), *q*_*variant,j*_ is the percent of the peptide variant *i* remaining intact after proteolysis by Saps, and *q*_*Hst*5,*j*_ is the percent of Hst5 remaining intact after proteolysis by Saps.

The *NIP* parameter sums the average amount of intact peptide remaining after incubation with each Sap individually and divides this value by the sum of the average amount of intact Hst5 remaining after incubation with each Sap individually. A value >1 indicates a variant is more proteolytically stable overall in the presence of multiple Saps than the parent Hst5, while values <1 indicates a variant is less proteolytically stable overall than Hst5. Based on the *NIP* for each peptide (**Table 2**), only K13R showed improved stability (*NIP* = 1.07) compared to Hst5, indicating a strongly positively charged residue at K13 contributes to stability and protection from proteolysis by Saps. The remaining K13 variants had *NIP* values ranging from 0.20 to 0.67, indicating an overall reduction in stability compared to Hst5. In contrast, substitutions at K17 led to an overall improvement in stability. The NIP values for the K17 variants ranged from 1.20 to 1.59, with K17W showing the most significant enhancement in overall proteolytic stability. Based on these results, K17 substitution should be considered as a starting point for designing peptides with improved proteolytic stability in environments with multiple Saps present.

**Table 2.** Normalized intact peptide values for Hst5 and K17 and K13 variants.

| Peptide | $NIP^a$ |
| --- | --- |
| Hst5 | 1.00 |
| K13R | 1.07 |
| K13L | 0.20 |
| K13H | 0.40 |
| K13Q | 0.24 |
| K13A | 0.51 |
| K13E | 0.67 |
| K17R | 1.20 |
| K17L | 1.35 |
| K17W | 1.59 |
| K17Q | 1.54 |
| K17A | 1.44 |
| K17E | 1.42 |

<sup>a</sup> *NIP* = normalized intact peptide

### Antifungal activity of full-length peptides

To determine the impact of amino acid substitutions at K13 and K17 on antifungal activity, we evaluated the antifungal activity of intact peptides against *C. albicans* (**Figure 3**). When assessing antifungal activity, we identified the lowest tested concentration where protease-treated Hst5 achieved over a 50% reduction in viability. We then compared the percent reduction in viability for each peptide at the identified concentration to determine statistical differences among variants. While the K13L, K13Q, K17L, K17R, and K17W variants had statistically similar antifungal activity to the parent Hst5 peptide, the K13A, K13E, K13Q, K13R, K17A, K17E, and K17Q variants showed a statistical difference in antifungal activity compared to Hst5. However, except for the K13E and K17E variants, these differences were modest, with all peptides surpassing a 50% reduction in viability by 4.7 µg/mL and a 90% reduction in viability by 9.4 µg/mL. When glutamate, a negatively charged amino acid, was substituted at K13 or K17, a 50% reduction was surpassed at 18.8 µg/mL and 9.4 µg/mL, respectively. Moreover a 95% reduction in viability was only surpassed at 75 µg/mL (the highest concentration tested). Our results underscore the negative impact of substituting a positively charged residue in Hst5 with a negatively charged residue, as these modifications can reduce antifungal activity.

**Figure 3.**
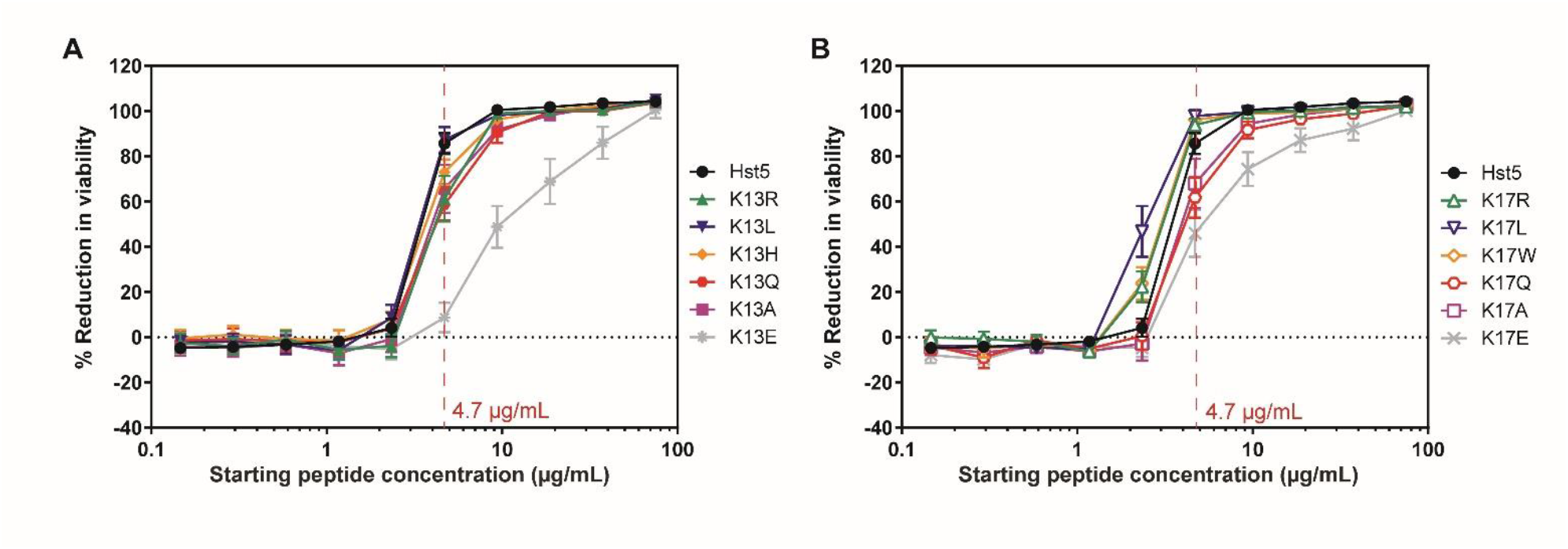
Antifungal activity of intact Hst5 variants with substitutions at A) K13 and B) K17. Hst5 variants were serially diluted and incubated with *C. albicans* cells for 30 min at 30 °C in 1 mM NaPB before quantifying the viability of the cells. The vertical dashed line and the written concentration denote the concentration at which statistical significance was assessed. Error bars represent the standard error of the mean (*n =* 6 for Hst5 and variants).

### Antifungal activity of K13 variants after treatment with Saps

In addition to testing the antifungal activity of the full-length peptide, we also assessed the antifungal activity after proteolysis by Saps. Degradation by many Saps, including Sap1, Sap2, Sap3, and Sap9, decreases the antifungal activity of the parent Hst5 peptide ^21-22, 28-29^, and proteolytic stability of Hst5 and variants of Hst5 is often correlated with retention in antifungal activity ^21-22, 29-30^. After incubation with Sap1 (**Figure 4A**), the antifungal activity of degraded Hst5 was modestly reduced compared to undegraded Hst5. All Sap1-degraded K13 variants had lower activity antifungal activity than degraded Hst5; however, both Hst5 and K13R surpassed a 90% reduction in viability at 4.7 µg/mL. After incubation with Sap2 (**Figure 4B**), the antifungal activity of Hst5 was noticeably reduced, and the antifungal activity of degraded K13R was similar to degraded Hst5. All other variants had reduced antifungal activity compared to Hst5 after incubation with Sap2. Incubation with Sap3 (**Figure 4C**) and Sap9 (**Figure 4D**) resulted in a substantial reduction in antifungal activity for Hst5, and all the K13 variants had even lower antifungal activity than degraded Hst5. Our results show that removing the positive charge residues at K13 can be inimical to the antifungal activity in the presence of Saps, illustrating the importance of the positive charge at K13.

**Figure 4.**
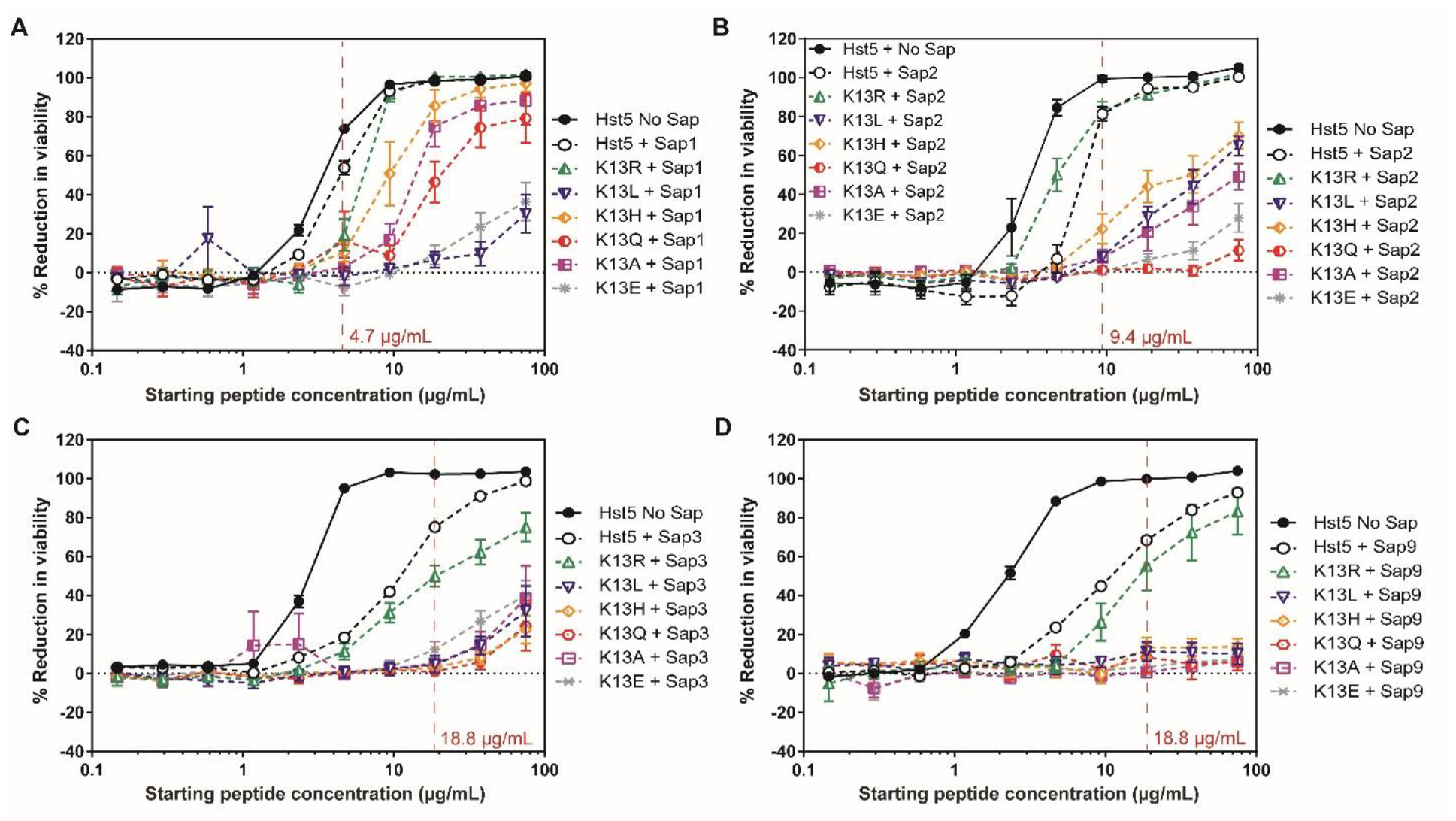
Residual antifungal activity of Hst5 and K13 variants after incubation with purified recombinant A) Sap1, B) Sap2, C) Sap3, or D) Sap9. Peptides were treated with Saps for 2 h at 37 °C and then incubated with *C. albicans* cells for 30 min at 30 °C in 1 mM NaPB before quantifying the viability of the cells. The vertical dashed line and the written concentration denote the concentration at which statistical significance was assessed. Error bars represent the standard error of the mean (for Sap1, Sap3, and Sap9, *n =* 12 for Hst5 controls with and without Sap and *n*=6 for variants; for Sap2, *n*=6 for Hst5 controls and variants).

### Antifungal activity of K17 variants after treatment with Saps

In contrast to the K13 variants, nearly all the degraded K17 variants retained higher levels of antifungal activity than the parent Hst5 peptide. After degradation by Sap1 (**Figure 5A**), the antifungal activity of degraded K17L was statistically similar to degraded Hst5, while all other K17 variants had statistically greater antifungal activity than degraded Hst5; however, the increase was quite modest, as all degraded variants and undegraded Hst5 surpass 50% reduction in viability at 2.3 µg/mL and surpass 90% at 4.7 µg/mL. With Sap2 (**Figure 5B**), proteolysis of K17E resulted in activity similar to that of degraded Hst5. However, the remaining degraded K17 variants had improved antifungal activity compared to the degraded Hst5. After incubation with Sap3 (**Figure 5C**), the only variant with reduced antifungal activity compared to Hst5 was K17L. In contrast, K17W was the only variant with a statistically significant improvement in antifungal activity compared to Hst5 following proteolysis by Sap3, with the reduction in viability surpassing 50% at 2.3 µg/mL. In comparison, the degraded Hst5 only surpassed 50% at 18.8 µg/mL. Finally, after incubation with Sap9 (**Figure 5D**), all K17 variants showed improved antifungal activity compared to degraded Hst5. Our results for the K17 variants highlight that substitutions at K17 generally lead to retained or enhanced antifungal activity after proteolysis by Saps; however, a leucine substitution can be greatly deleterious in the presence of Sap3.

**Figure 5.**
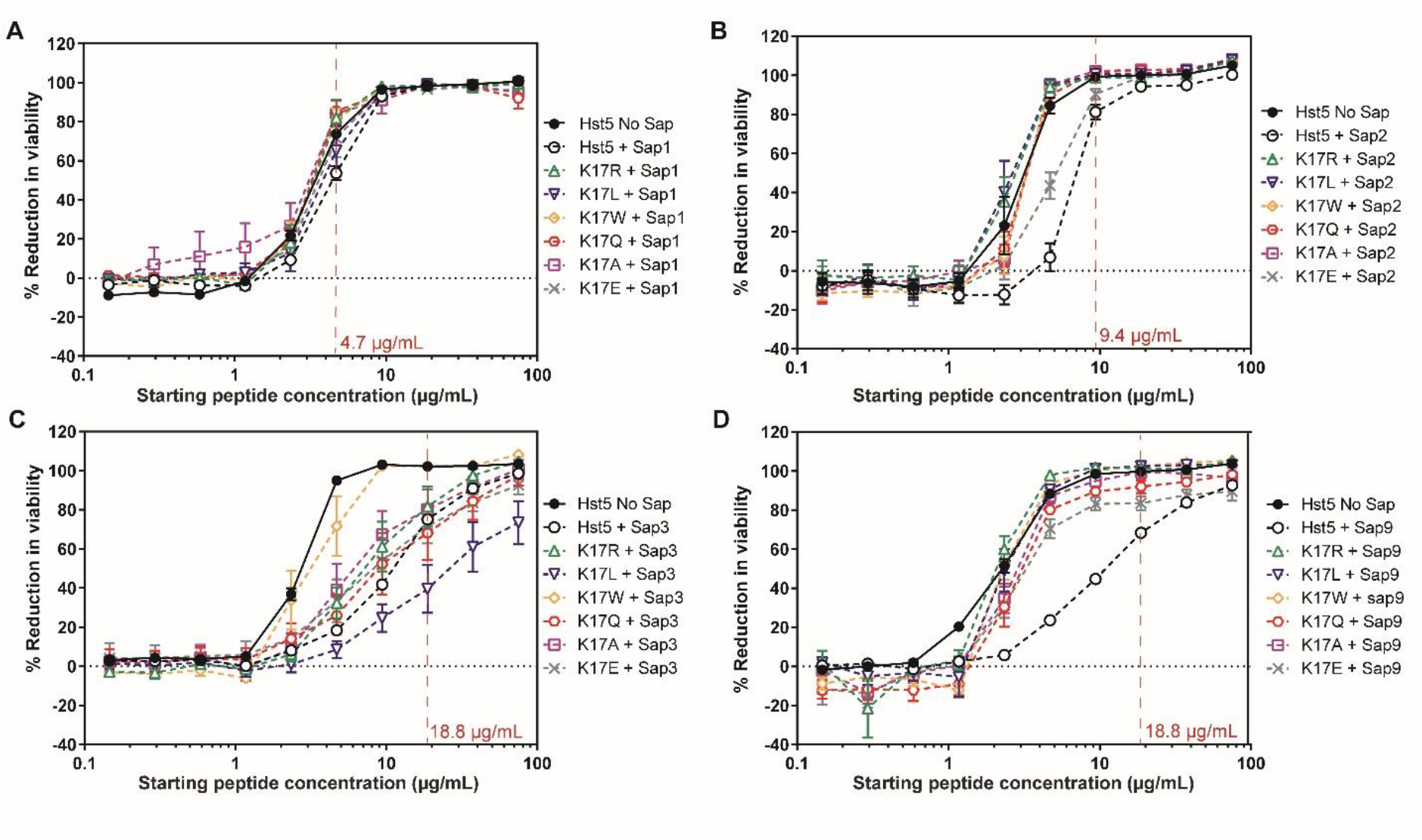
Residual antifungal activity of Hst5 and K17 variants after incubation with purified recombinant A) Sap1, B) Sap2, C) Sap3, or D) Sap9. Peptides were treated with Saps for 2 h at 37 °C and then incubated with *C. albicans* cells for 30 min at 30 °C in 1 mM NaPB before quantifying the viability of the cells. The vertical dashed line and the written concentration denote the concentration at which statistical significance was assessed. Error bars represent the standard error of the mean (for Sap1, Sap3, and Sap9, *n =* 12 for Hst5 controls with and without Sap and *n*=6 for variants; for Sap2, *n*=6 for Hst5 controls and variants).

### Proteolytic stability of saliva-treated Hst5 variants

To understand how amino acid substitutions impact the degradation of Hst5 and Hst5 variants by saliva, we incubated each Hst5 variant with saliva and quantified the degradation, as we did for the Saps. After incubating Hst5 with saliva (**Figure 6**), 37.7% of Hst5 remained intact. Most of the K13 variants were as stable as the parent Hst5 in saliva, though K13R exhibited decreased stability (29.2 % remaining intact) (**Figure 6A**). After incubation of the K17 variants with saliva (**Figure 6B**), K17L and K17W showed an improvement in proteolytic stability, with 52.9% and 52.8% of the peptides remaining intact, respectively. The remaining K17 variants were degraded similarly to the parent peptide. The improvement in the stability of K17L and K17W could be related to the hydrophobicity and large size of the leucine and tryptophan side chains. The modest improvement in proteolytic stability of K17L and K17W with saliva compared to the improvements seen for the same variants with the Saps indicates that engineering proteolytic stability against Saps alone is not sufficient to substantially protect against other proteases in saliva. However, our variants with improved stability in saliva provide a reasonable starting point for designing more broadly proteolytically stable variants.

**Figure 6.**
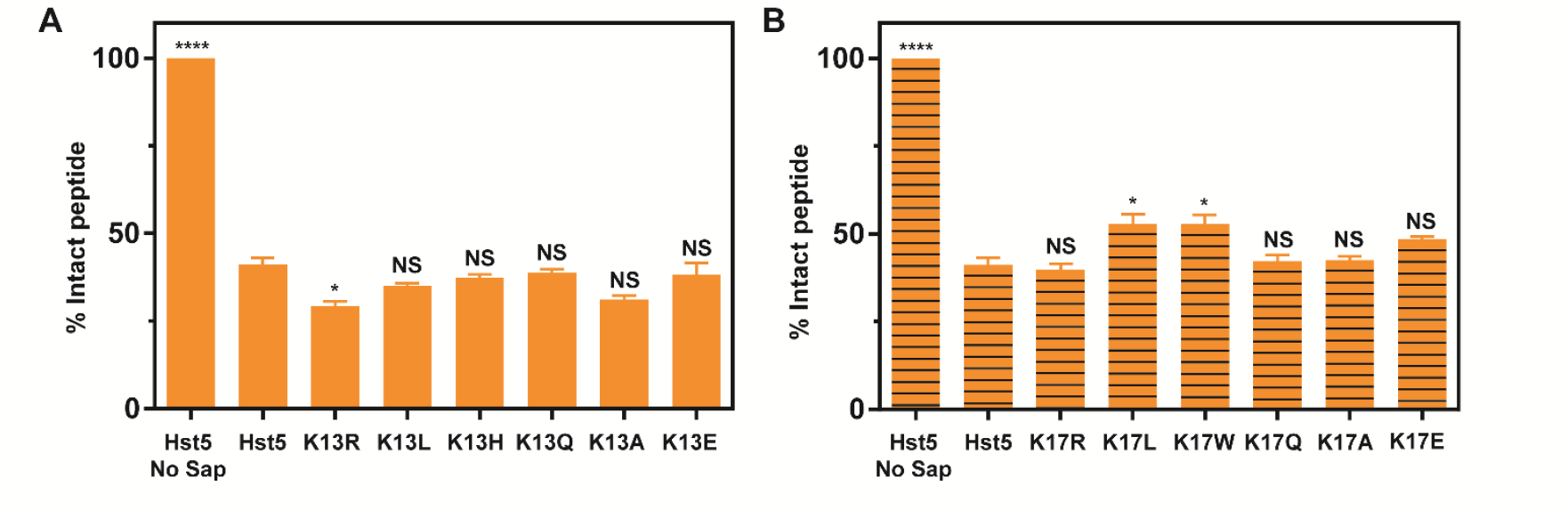
Degradation of Hst5 and A) K13 and B) K17 variants following incubation with saliva. Peptides were incubated with saliva for 2 h at 37 °C, and intact peptides and their degradation fragments were separated by gel electrophoresis. The percentage of peptide remaining intact was quantified by densitometry. Error bars represent the standard error of the mean (*n =* 6 for the Hst5 controls with and without saliva and *n =* 3 for the Hst5 variants). Representative gel images are provided in **Figure S5**.Statistical significance, relative to degraded Hst5, is indicated as follows: ns for not significant, ns for not significant, * for p ≤ 0.05, ** for *p* ≤ 0.01, *** for *p* ≤ 0.005, and **** for *p* ≤ 0.0001.

Since K17W was found to be the most proteolytically stable variant after incubation with Saps and saliva, we further analyzed the fragments formed by the saliva degradation using mass spectrometry (**Figure S6**). These data show that K17W enhances proteolytic stability specifically around the K17 site, resulting in 52% greater retention of the full-length peptide compared to Hst5; however, residues between K11 and F14 remain susceptible to cleavage. The results suggest that pairing K17W with additional substitutions across the K11–F14 region may provide a strategy to further increase proteolytic stability.

### Antifungal activity of saliva-treated Hst5 variants

In addition to assessing the proteolytic stability of Hst5 variants in saliva, we evaluated the antifungal activity of the fragments produced from incubation with saliva. After degradation by saliva, the antifungal activity of Hst5 was reduced (**Figure 7**). Of the K13 variants, only degraded K13R retained similar activity to degraded Hst5, while all other K13 variants showed reduced activity (**Figure 7A**). Similarly, all K17 variants exhibited diminished activity compared to Hst5 after incubation with saliva, except K17R and K17W (**Figure 7B**). Degraded K17R exhibited comparable antifungal activity to degraded Hst5, and K17W was the only variant with enhanced antifungal activity compared to Hst5 following incubation with saliva. Our findings suggest that the positive charge at K13 and K17 may generally help the parent Hst5 peptide retain its antifungal activity in the oral cavity. However, introducing tryptophan at K17 can enhance the antifungal activity following exposure to saliva, which may be related to its modest improved resistance to proteolysis by saliva.

**Figure 7.**
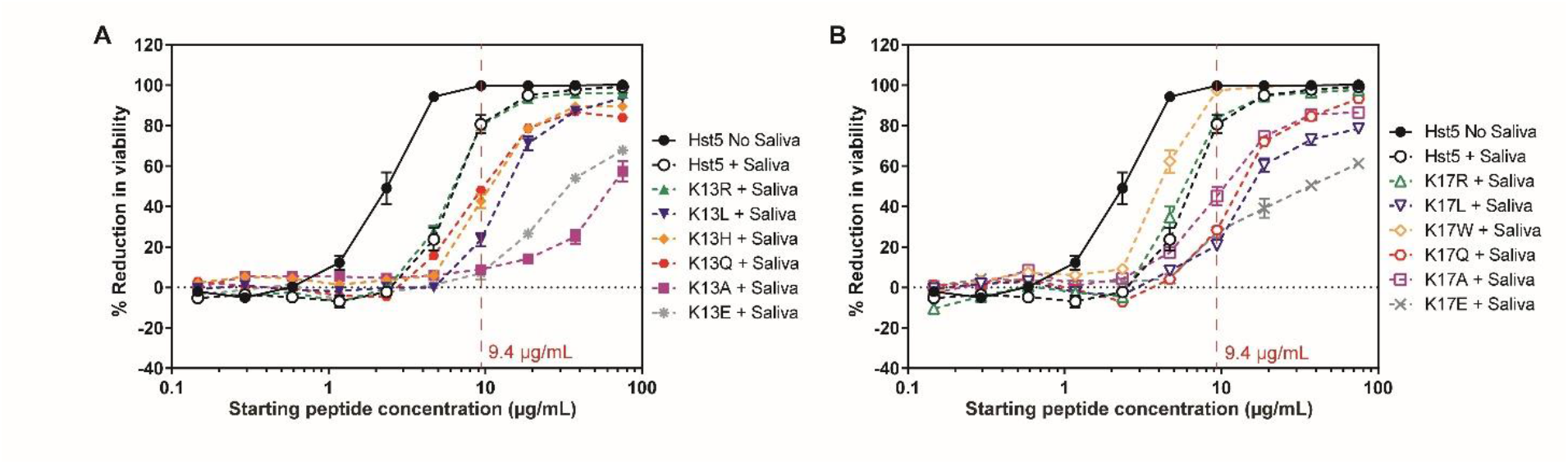
Residual antifungal activity of Hst5 and A) K13 and B) K17 variants after incubation with saliva. Peptides were treated with saliva for 2 h at 37 °C and then incubated with *C. albicans* cells for 30 min at 30 °C in 1 mM NaPB before quantifying the viability of the cells. The vertical dashed line and the written concentration denote the concentration at which statistical significance was assessed. Error bars represent the standard error of the mean (*n =* 6 for Hst5 controls and variants).

### Biofilm inhibition ability of Hst5 and K17W

Having designed a Hst5 variant with improved proteolytic stability and antifungal activity against the yeast form of *C. albicans*, we next assess the ability of the K17W variant to prevent biofilm growth. Using a metabolic assay with (2,3-bis-(2-methoxy-4-nitro-5-sulfophenyl)-2*H*-tetrazolium-5-carboxanilide) (XTT), we quantified biofilm formation following incubation with Hst5 and K17W across the tested concentration range (**Figure 8**). At the highest concentration (750 µM), Hst5 reduced biofilm formation by 39.1% while K17W achieved an 84.1% reduction. At 188 µM, Hst5 did not inhibit biofilm growth, while K17W reduced growth by 59.2%. Altogether, these results demonstrate that engineering proteolytically stable variants of Hst5 can improve both antifungal and antibiofilm efficacy, with K17W exhibiting the strongest performance among the tested variants.

**Figure 8.**
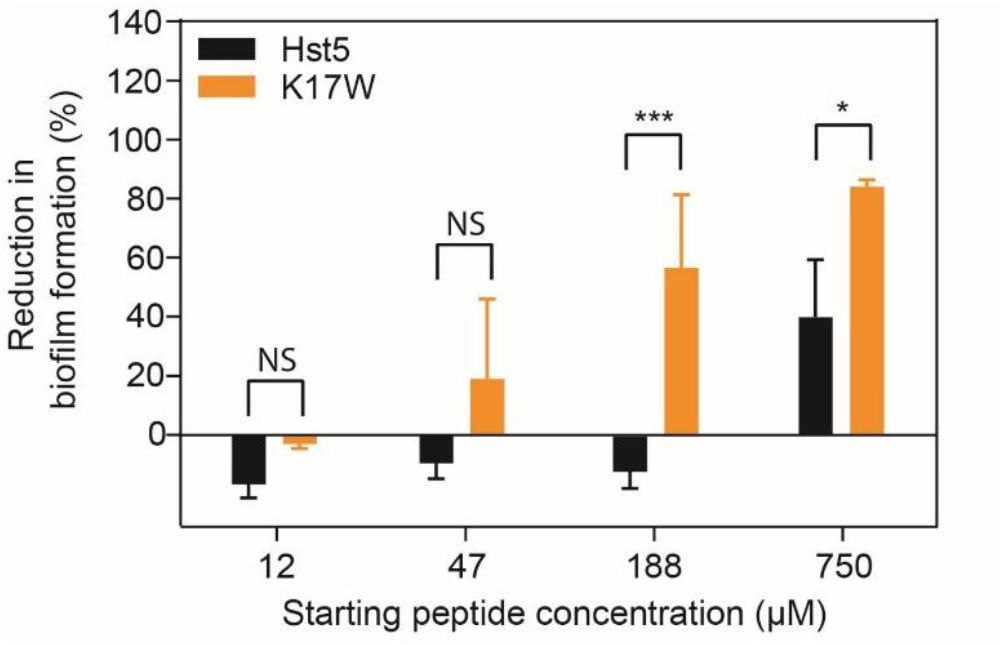
Prevention of biofilm formation by Hst5 and K17W. Peptides were serially diluted and incubated with *C. albicans* cells for 24 h at 37 °C in RPMI1640 medium. Reduction of biofilm formation at select concentrations was measured using an XTT-based. Error bars represent the standard error of the mean (*n* = 4). Statistically significant differences, relative to degraded Hst5, are indicated as follows: ns for not significant, * for *p* ≤ 0.05, and *** for *p* ≤ 0.005.

## Discussion

In this work, we examined how modification of Hst5 at K13 and K17 affects interactions with Saps and saliva. Of the modifications we studied, five were introduced previously by our lab (K13R, K13L, K13H, K17R, and K17L) ^21-22, 30^, one was introduced by Tsai et al. (K13E) ^31^, and six were new to this work (K13A, K13Q, K17A, K17E, K17Q, and K17W). We assessed the proteolytic stability and antifungal activity of these peptide variants both before and after incubation with Saps and saliva determined how the modifications influence Hst5’s proteolytic stability and antifungal efficacy.

### Retaining a positive charge at K13 is important for proteolytic stability and antifungal activity

Our results highlight the benefits of retaining the positive charge at K13. For all the Saps tested, the K13R modification maintained or improved the proteolytic stability compared to Hst5, indicating that the positive charge at K13 is likely important in maintaining proteolytic stability of Hst5 and Hst5 variants in the presence of Sap1, Sap2, Sap3, and Sap9. In contrast, reducing the positive charge (K13H) or removing the positive charge (K13L, K13Q, K13A, K13E) led to increased proteolysis in nearly all cases. The exceptions to this are K13E with Sap3 and K13A with Sap9. The retention of proteolytic stability by K13E after incubation with Sap3 aligns with prior work by Koelsch et al. that suggested Sap3 has a reduced affinity for the glutamate side chain compared to Sap1 and Sap2 ^20^. In terms of antifungal activity, all K13 variants except for K13R experienced a significant reduction in activity after incubation with Saps, which can likely be linked to the reduced proteolytic stability. Although K13R (after incubation with Sap3 and Sap9), K13E (after incubation with Sap3) and K13A (after incubation with Sap9) exhibited similar proteolytic stability to Hst5, their reduced antifungal activity compared to Hst5 indicates that the fragments formed have shifted to fragments with reduced antifungal activity, which is consistent with previous work that showed the identity of the fragments produced by proteolysis can impact antifungal activity ^21-22, 30^. Overall, our results emphasize that, while variants with substitutions at K13 can retain proteolytic stability, retaining the positive charge at K13 is crucial for preserving antifungal efficacy.

### Modifications at K17 generally improve proteolytic stability and antifungal activity

In contrast to the results for the K13 variants, modifications at K17 were often associated with improved proteolytic stability and antifungal activity. In general, K17 variants were as stable or more stable than the parent Hst5 in the presence of Sap 1, Sap3, and Sap9, consistent with previous work from our lab demonstrating that the K17R and K17L substitutions enhance proteolytic stability during incubation with Sap9, *C. albicans* cells, and *C. albicans* culture supernatant ^21, 30^. Our findings suggest that modifications to K17 can generally maintain or improve protection of Hst5 from proteolysis by Saps. Additionally, the only variants with improved stability in the presence of saliva were K17 variants (K17L and K17W). Similarly, the antifungal activity of nearly all K17 variants after incubation with Saps (the exception being K17L treated with Sap3) was either the same or greater than the parent Hst5 peptide. Combined, our findings suggest a link between the proteolytic stability and antifungal activity of the K17 variants and emphasize that K17 should be prioritized in designing variants for increased stability without compromising activity.

### Glutamate substitution for K13 and K17 in Hst5 should be avoided

To exert its activity, Hst5 must interact with on the surface of *C. albicans* cells ^10^. We and others have previously observed that the antifungal activity of Hst5 is generally tolerant to arginine or leucine substitutions at its four lysine residues ^21-22^, though the K13E and K13T substitutions can reduce antifungal activity ^30-31^. Our current results show that both the K13E and K17E substitutions resulted in a reduction in antifungal activity, though substitutions to amino acids with neutral side chains at K13 and K17 did not greatly impact antifungal activity. Previous work from our lab showed E16R and E16L substitutions improved antifungal activity ^30^, further highlighting that negatively charged glutamate residue may reduce Hst5’s ability to interact with the surface of *C. albicans* cells and is undesirable for strong Hst5 antifungal activity. Variants with substitutions to glutamate also do not offer significant advantages compared to other Hst5 modifications in terms of the proteolytic stability in the presence of Saps and saliva, suggesting that these residues should be avoided when designing improved Hst5 variants.

### NIP parameter reveals K17W as the most impactful substitution

Prior to this work, studies on the proteolytic stability of Hst5 and its variants in the presence of Saps assessed cleavage by individual Saps without a quantitative analysis that incorporated the overall impact of multiple Saps ^21-22, 29^. However, multiple Saps are simultaneously expressed by *C. albicans*, including during infection ^32-34^, so having a quantitative method to incorporate the effect of multiple Saps on Hst5 degradation is desirable. We developed the NIP parameter to provide this quantitative analysis and allow us to determine which Hst5 modifications provide the best overall stability in the presence of multiple Saps. Our results revealed that nearly all K13 variants had worse overall proteolytic stability than Hst5, though K13R had a small improvement in stability. NIP also reveals differences in the overall stability of the K17 variants. Despite all variants providing an overall improvement in proteolytic stability, the K17R variant that retained the positive charge at K17 showed the least improvement and the K17W variant showed the greatest improvement in proteolytic stability. Based on these results, combining K13R with targeted K17 modifications, such as K17W, offers a promising strategy to enhance proteolytic stability and retain proteolytic stability after incubation with Saps. The version of the NIP equation that we employed in our analysis only incorporates Sap1, Sap2, Sap3, and Sap9, since these were the Saps relevant in this work. However, studies with other peptides could incorporate additional Saps or other relevant enzymes to tailor NIP for other applications. Moreover, the NIP parameter could be modified to provide different weights to each enzyme to tailor the parameter for a specific environment or disease condition where enzymes have differential levels of importance. The NIP parameter will be a useful tool to identify the peptide modifications that lead to improved overall resistance to proteolysis when the peptide will be used in an environment with multiple enzymes.

### K17W variant shows enhanced proteolytic stability and antifungal activity after incubation with saliva

As a salivary peptide and proposed therapeutic for oral candidiasis, Hst5 may interact with the complex mixture of host and microbial enzymes in saliva. Prior work showed that saliva can fragment Hst5 ^7, 25, 27, 35-36^, and we evaluated how the variants of Hst5 performed after incubation with saliva K13 variants exhibited proteolytic stability in saliva that was comparable to the parent Hst5, except for K13R, which was degraded more than Hst5. In contrast, the K17L and K17W variants had modest improvements in proteolytic stability in saliva compared to the parent peptide, which may be due to their bulky, hydrophobic side chains reducing interactions with enzymes in saliva. Our proteolysis results indicate that, while *C. albicans* Saps contribute to the mixture of enzymes in saliva, Saps are likely not the primary drivers of Hst5 proteolysis by saliva. Interestingly, when examining antifungal activity post-saliva degradation, only the K17W variant had enhanced activity compared to saliva-degraded Hst5. Variants with arginine substitutions (K13R and K17R) had similar activity to Hst5, while all other variants had reduced stability. Based on these results, K17W offers the greatest protection from both Saps and salivary proteases, as well as the most significant enhancements in antifungal activity following incubation with Saps and saliva, making it a compelling starting point in the design of future Hst5 variants.

### K17W variant prevents biofilm formation

Biofilm formation involves the formation of dense networks of cells and can significantly enhance resistance to treatment with standard antifungal agents ^37-40^. Prior work has shown that proteolytically stable variants of Hst5 can improve antibiofilm efficacy ^37-41^, so we evaluated the ability of our most stable variant (K17W) to prevent biofilm formation. Although complete inhibition of biofilm formation was not observed, K17W surpasses a 50% reduction in biofilm viability at concentrations above 188 µM, whereas Hst5 never surpasses 50% within the concentration range tested. Achieving a comparable reduction in biofilm viability required 50-fold more peptide than was needed to inhibit the growth of yeast cells, but this difference is smaller than the 1000-fold increase in resistance observed for some small-molecule therapeutics^39-40, 42^. Our findings indicate that rationally engineered Hst5 variants can meaningfully reduce biofilm formation on surfaces, even within the challenging context of biofilm-associated resistance.

## Conclusion

Studying variants of Hst5 with modifications at K13 and K17 helped elucidate valuable information about the role these residues play in interactions between Hst5 and salivary and fungal proteases. When the positive charge at K13 is removed, fragments resulting from the incubation of the variants with Saps and saliva had reduced antifungal activity compared to Hst5, showing the importance of retaining the positive charge at K13. In contrast, variants with modifications at K17 generally showed an improvement in proteolytic stability in the presence of Saps and retained antifungal activity following incubation with Saps. By introducing the NIP parameter, we identified the K17W variants as the most likely to remain stable in environments containing multiple Saps. The K13 and K17 Hst5 variants all remained susceptible to proteolysis by saliva, but the K17W variant showed greater stability and retained more antifungal activity than the other variants and showed enhanced biofilm prevention compared to the parent Hst5 peptide. Our results with both Saps and saliva suggest the K17W variant as a starting point for further work in designing Hst5 for proteolytic stability and antifungal activity. The strategies and techniques we used provide valuable insights into the relationship between proteolytic stability and antifungal activity and could be applied to designing a broad range of antimicrobial peptides to improve their therapeutic potential.

## Experimental section

### Preparation of peptides and proteases

The parent Hst5 peptide and Hst5 variants were commercially synthesized by Biomatik (Ontario, Canada) with a purity ≥95% and trifluoroacetic acid removal to hydrochloric acid salt. The peptides were provided in a lyophilized form and reconstituted in water at a concentration of 4 mg/mL to create stock solutions for use in assays. Purified recombinant Sap1, Sap2, Sap3, Sap5, Sap6, Sap9, and Sap10 were prepared as described previously ^14^. Briefly, Sap1, Sap2, Sap3, and Sap6 were purified using ion exchange chromatography and desalted into 0.1 M sodium citrate buffer. Sap5 was purified by ultrafiltration and desalted into 0.1 M sodium citrate. Sap9 and Sap10 were expressed without a GPI anchor and then purified by ion exchange chromatography and desalted into 0.1 M sodium citrate buffer.

### Assay of proteolytic degradation of peptides by Saps

The extent of degradation of the peptides by the Saps was assessed by incubating each Sap with each peptide variant. The Sap concentrations were selected so ∼50% of the parent peptide Hst5 was degraded under the assay conditions: 5.0 μg/mL for Sap1, 0.1 μg/mL for Sap2, 1.6 μg/mL for Sap3, 5.0 μg/mL for Sap5, 5.0 μg/mL for Sap6, 3.1 μg/mL for Sap9, and 5.0 μg/mL for Sap 10. The Saps were individually incubated with each Hst5 variant (150 μg/mL) in 1 mM sodium phosphate buffer (NaPB) at a pH of 7.4 for 2 h at 37 °C. After incubation, samples were boiled at 100 °C for 5 minutes to inactivate the proteases and stored at -20 °C. No degradation was observed under the same conditions without the Saps present.

### Assay of proteolytic degradation of peptides by saliva

Frozen, unfiltered, gender-pooled saliva was sourced from BIOIVT (Westbury, NY). Cells and debris were removed from the saliva by centrifugation at 3,900 ×g for 5 minutes. The resulting supernatant was passed through a 10 kDa molecular weight cutoff column to exchange the buffer to 2 mM NaPB. The processed saliva was then diluted in 2 mM NaPB to a concentration of 1.0 mg/mL total protein concentration and stored at −20 °C until subsequent use. The degradation of the peptides by saliva was evaluated using a strategy analogous to that used for degradation by Saps. The peptides (150 μg/mL) and saliva (500 μg/mL total protein) were incubated in 1 mM NaPB at 37 °C for 2 h, boiled for 5 min at 100 °C to inactivate the proteases in saliva, and stored at 20 °C until analyzed.

### Analysis of peptide degradation by gel electrophoresis

Following proteolysis by either Saps or saliva, each peptide sample was thawed, mixed with tricine samples buffer (200mM Tris-HCl, pH 6.8, 40% glycerol, 2% sodium dodecyl sulfate), and supplemented with 2% β-mercaptoethanol. The mixture was boiled for 5 min at 100 °C. Full-length peptides and degraded fragments were separated by gel electrophoresis using a 16.5% Tris-tricine gel (Bio-Rad, Hercules, CA). After electrophoresis, gels were fixed in a solution containing 10% acetic acid, 40% methanol, and 50% water for 30 minutes. Fixed gels were stained using Bio-Safe Coomassie stain (Bio-Rad) for 1 hour and washed in ultrapure water. The gels were imaged using a ChemiDoc imager (Bio-Rad), and densitometric analysis was conducted with the Image Lab software (Bio-Rad) to determine the percentage of the peptide remaining intact. The intact peptide was identified as the upper band in each lane at ∼3000 Da, and all bands below this were considered degradation products.

Statistical analysis of the gel data was performed using two-way ANOVA tests with *α* = 0.05 and Dunnet’s multiple comparison tests, using degraded Hst5 as the control. Representative gel images are provided in **Figure S1, S2, S3, S4, and S5**, and the corresponding *p*-values for the comparison tests are summarized in **Table S1**.

### Assay of antifungal activity of full-length peptide

The antifungal activity of intact peptide variants was assayed using an optical density-based candidacidal assay, as previously described ^43^. *C. albicans* strain SC5413 cells were inoculated into yeast extract-peptone-dextrose (YPD) medium (20 g/L dextrose, 20 g/L peptone, 10 g/L yeast extract) and incubated overnight at 30 °C while shaking at 230 RPM. Cell were grown at 30 °C to promote yeast cell growth and limit hyphal growth ^44^. After the overnight incubation, cells were subcultured to an optical density at 600 nm (OD_600_) of 0.1 and grown at 30 °C until reaching an OD_600_ of ∼1.0. Cells were washed and diluted to a cell density of 5.0 × 10^5^ cell/mL in 2mM NaPB. Separately, 20 µL serial dilutions of peptide variants were prepared in water in a 96-well plate. The prepared cell suspension was added to each well, resulting in final concentrations of 0.14 – 75 µg/mL peptide and 2.5 × 10^5^ cells/mL in 1 mM NaPB. After incubation of the peptide-cell mixtures at 30 °C for 30 minutes, 280 µL of 1 mM NaPB was added to each well to reduce interactions between peptide and cells. From each well, a volume of 8 µL (containing ∼250 cells) was inoculated into a new flat bottom 96-well plate containing 100 µL NaPB and 100 µL YPD. Wells containing YPD and NaPB without cells served as a negative control. The new plate was incubated overnight at 30 °C while shaking at 350 RPM. After incubation, cells in each well were resuspended, and the OD_600_ was measured. The percentage reduction in viability was then calculated from

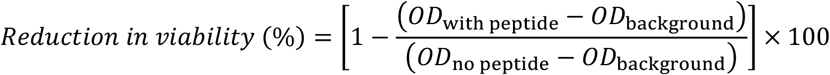

where *OD*_with peptide_ is the OD_600_ of a well containing cells and peptide, *OD*_no peptide_ is the OD_600_ of a well containing cells with no peptide, and *OD*_background_ is the average OD_600_ of wells containing the negative control with NaPB and YPD only. The assay was performed on three separate days with two replicates on each day. To perform a statistical comparison of the data, the lowest tested concentration of the protease-treated parent Hst5 peptide that exceeded 50% reduction in viability was identified. The reduction of viability for each peptide at this concentration was then compared using two-way ANOVA tests with α = 0.05 and Dunnett’s multiple comparison tests (degraded Hst5 as the control). *P* < 0.05 was deemed as significant (**Table S2**).

### Assay of antifungal activity of Sap- and saliva-treated peptides

To assay the antifungal activity of the peptides following degradation by Saps or saliva, we utilized a method similar to that for testing the antifungal activity the intact peptides, but the peptides were first incubated with the Saps or saliva. For Sap-treated peptide, the degradation was performed in the same manner as described above for the proteolytic degradation assay. For saliva-treated peptide, the assay was performed as described for the proteolytic degradation assay, except the saliva was passed through a 0.2 µm syringe filter to ensure sterility, and the saliva concentration in the assay was reduced to a final total protein concentration of 115 µg/mL to better visualize differences in activity between saliva-treated Hst5 and variants. Serial dilutions (20 µL) of degraded peptide variants were then prepared in water in a 96-well plate.

A suspension of *C. albicans* cells was prepared as described for the assay with intact peptides, except the cell suspension was prepared in 1 mM NaPB. The cell suspension was added to the serial dilutions of degraded peptide, resulting in wells containing the equivalent of 0.14 – 75 µg/mL peptide (based on the peptide concentration prior to degradation) and 2.5 × 10^5^ cell/mL in 1 mM NaPB. After incubation of the degraded peptides with cells at 30 °C for 30 minutes, the samples were processed as described for the assay with intact peptides, and the OD_600_ of was measured to calculate the percent reduction in viability. As with the intact peptide assay, statistical analysis was done at the lowest tested concentration where the reduction in viability due to degraded Hst5 exceeded 50%. Two-way ANOVA tests with α = 0.05 and Dunnett’s multiple comparison tests were performed to compare the degradation product for each peptide to the degradation products for Hst5. *p*-values are provided in **Table S2**.

### Assay of biofilm prevention by full-length peptides

The ability of the peptide to prevent biofilm formation was assessed as we have done previously reported^41, 45^. A single colony of *C. albicans* SC5314 was inoculated into YPD and incubated overnight at 30 °C. After overnight growth, cells were subcultured to an OD_600_ of 0.1 and grown at 30 °C until reaching an OD_600_ of ∼1.0. Cells were washed in 1× phosphate-buffered saline (PBS) and diluted to an OD_600_ of 1×10^6^ cells/mL in RPMI 1640 medium (with L-glutamine, and without sodium bicarbonate) (MP Biomedicals; Santa Ana, California) buffered with MOPS (3-[N-morpholino] propanesulfonic acid) (Gibco; Waltham, MA) at a pH of 7.0. Separately, 50 μL serial dilutions of peptides were prepared in water in a 96-well plate. The prepared cell suspension was added to each well, resulting in a final concentration of 750 μg/mL peptide and 5×10^5^ cells/mL in 0.5× RPMI 1640. The plate was incubated at 37 °C for 24 h to allow biofilm growth ^44^. After incubation, the plates were decanted, and the wells were washed with 1× PBS. The wash buffer was removed, and 100 μL of an XTT working solution (0.5 g/L XTT and 1 μM menadione) was added to each well. The plate was then incubated for 90 minutes in the dark. After incubation, 75 μL of the solution was transferred to a corresponding well in a new 96-well plate, and the absorbance was measured at 490 nm using a plate reader. The percent reduction in biofilm formation was calculated as

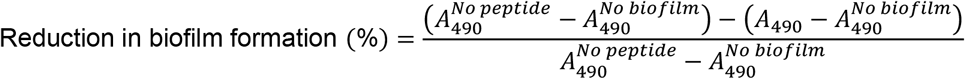

where 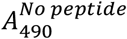 is the average absorbance for wells containing cells with no peptide,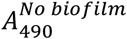 is the average absorbance for wells containing RPMI 1640 and no cells, and *A*_490_ is the absorbance for wells containing both cells and peptide. This assay was performed with two replicates on two separate days. To perform statistical analysis, a two-way ANOVA test was used with α = .05 and Dunnett’s multiple comparison tests (Hst5 as the control) at each concentration tested (**Table S3**).

### Identification of peptide fragments using mass spectrometry

Mass spectrometry was used to identify and quantify peptide fragments generated after incubation with Saps or whole saliva. Following degradation assays, 30 μL of each sample was desalted using C18 TopTip columns (Glygen, Columbia, MD) with 0.1% formic acid for binding and 0.1% formic acid/80% acetonitrile for elution. A 19 μL aliquot was mixed with 1 μL of the peptide MRFA (0.01 μg/mL) as an internal standard. Samples were directly injected into a Bruker Maxis II mass spectrometer (Billerica, MA), acquiring data from *m*/*z* 250–3200 at >80,000 FWHM. Single replicates were collected for each peptide–saliva condition. Spectra were processed using Bruker Compass DataAnalysis and BioTools. Deconvoluted spectra were matched to theoretical fragment masses (±0.2 *m*/*z*). Fragments longer than seven residues were retained, and the fragments with normalized intensities >10% of the highest-intensity fragments were included in the final analysis.

## Supporting information

This article contains supporting information (Makambi et al. - Supporting information.pdf). This file includes representative gel images for degradation by all Saps and statistical analysis data.

## Author contributions

WKM: Conceptualization, formal analysis, investigation, methodology, project administration, validation, visualization, writing – original draft preparation, writing – review & editing. VLC: Investigation, writing – review & editing. LK: Resources, review & editing. BH: Resources, review & editing. AJK: Conceptualization, funding acquisition, methodology, project administration, supervision, validation, visualization, writing – review & editing.

## Acknowledgments

We thank Stephanie Wisgott (Leibniz-HKI) for technical support in purifying the Sap enzymes. This work was supported by the National Institutes of Health (R03DE029270 to AJK), and WKM was supported by a Department of Education fellowship (GAANN-P200A180093). The content is solely the responsibility of the authors and does not necessarily represent the official views of the National Institutes of Health.

## Abbreviations

Hst5: histatin 5
Sap: secreted aspartyl proteases
GPI: glycosylphosphatidylinositol
MOPS: 3-(N-morpholino) propanesulfonic acid
NaPB: sodium phosphate buffer
NIP: normalized intact peptide
OD: optical density
PBS: phosphate-buffered saline
XTT: (2,3-bis-(2-methoxy-4-nitro-5-sulfophenyl)-2*H*-tetrazolium-5-carboxanilide)
YPD: yeast extract-peptone-dextrose

