## Supplementary figures and tables for "Optimizing histatin 5: Effects of K13 and K17 substitutions on proteolytic stability and antifungal activity"

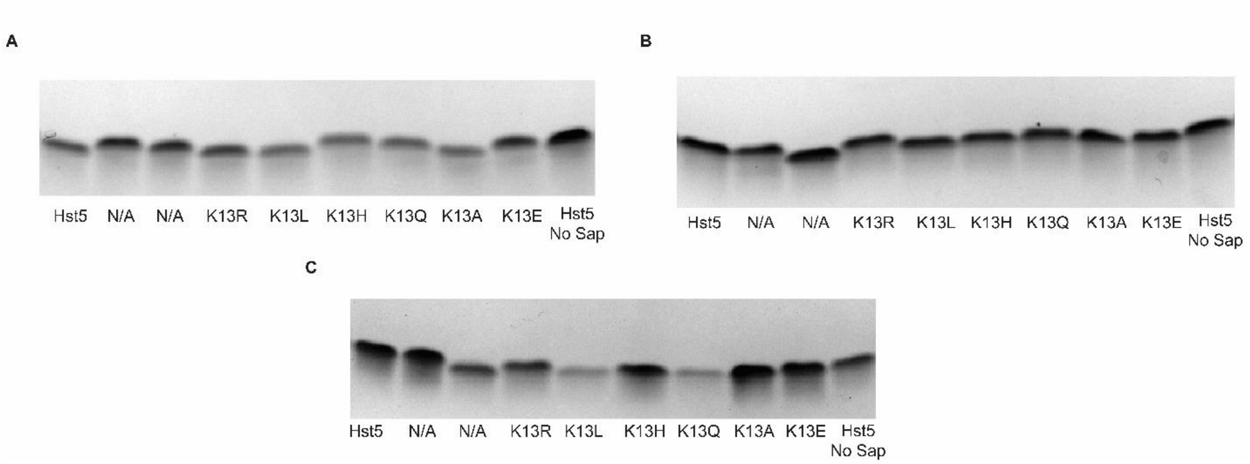

**Figure S1.** Representative gel images for proteolysis of Hst5 and variants with modifications at K13 by recombinant (A) Sap5, (B) Sap6, and (C) Sap10. Lanes marked "N/A" contain variants that are not relevant to this work.

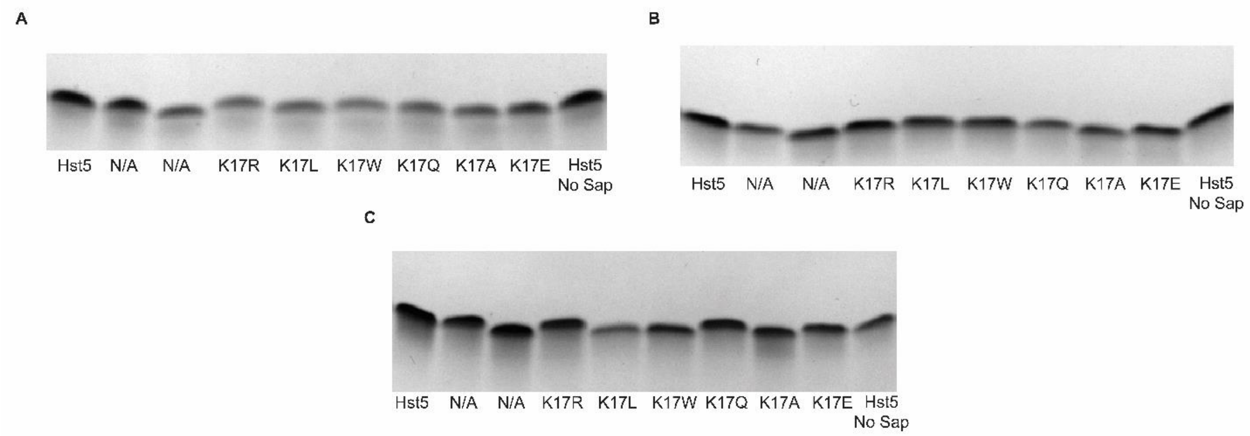

**Figure S2.** Representative gel images for proteolysis of Hst5 and variants with modifications at K17 by recombinant (A) Sap5, (B) Sap6, and (C) Sap10. Lanes marked "N/A" contain variants that are not relevant to this work.

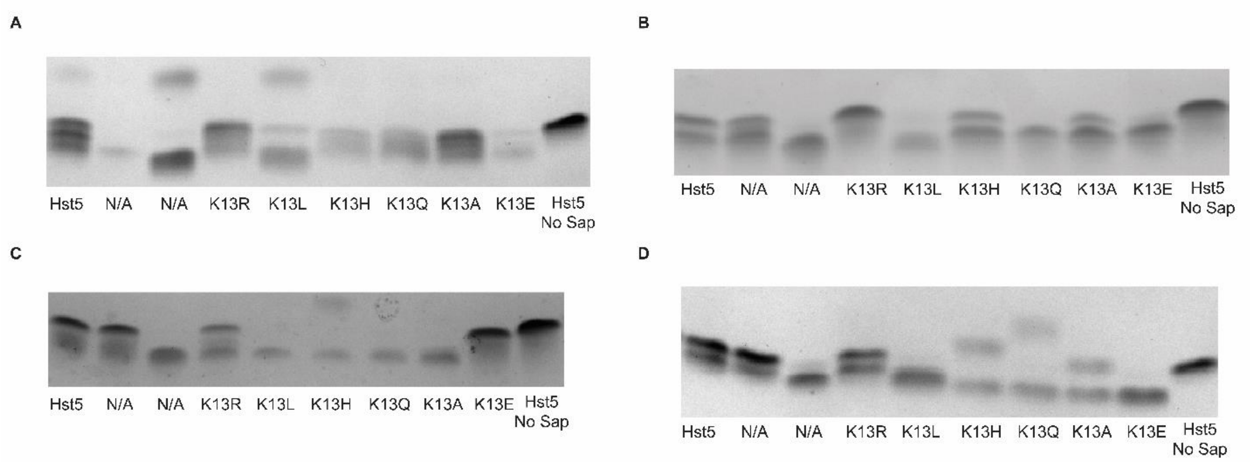

**Figure S3.** Representative gel images for proteolysis of Hst5 and variants with modifications at K13 by recombinant (A) Sap1, (B) Sap2, (C) Sap3, and (D) Sap9 Lanes marked “N/A” contain variants that are not relevant to this work.

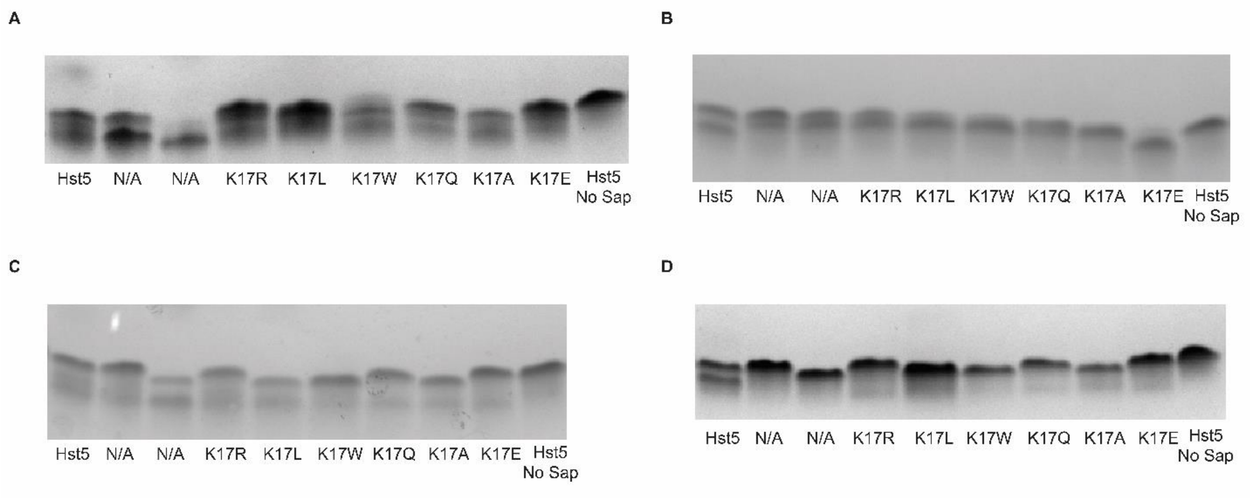

**Figure S4.** Representative gel images for proteolysis of Hst5 and variants with modifications at K17 by recombinant (A) Sap1, (B) Sap2, (C) Sap3, and (D) Sap9. Lanes marked “N/A” contain variants that are not relevant to this work.

**Table S1.** *p*-Values for data evaluating proteolysis of Hst5 and variants by Saps and saliva (Figures 1, 2, and 6).

| Peptide | *P*-values (Proteolysis assay)[a] | | | | |
|---|---|---|---|---|---|
| | Sap1 | Sap2 | Sap3 | Sap9 | Saliva |
| K13R | 0.9145 | 0.0286 | 0.9998 | 0.9877 | 0.0122 |
| K13L | < 0.0001 | < 0.0001 | < 0.0001 | < 0.0001 | 0.4414 |
| K13H | 0.0016 | < 0.0001 | < 0.0001 | < 0.0001 | 0.8503 |
| K13Q | 0.0001 | < 0.0001 | < 0.0001 | < 0.0001 | 0.9854 |
| K13A | 0.0079 | < 0.0001 | < 0.0001 | 0.946 | 0.0507 |
| K13E | 0.0013 | < 0.0001 | 0.0008 | < 0.0001 | 0.9614 |
| K17R | 0.059 | 0.010 | 1.000 | < 0.0001 | 1.000 |
| K17L | 0.160 | < 0.0001 | 0.940 | < 0.0001 | 0.047 |
| K17W | 0.082 | < 0.0001 | < 0.0001 | < 0.0001 | 0.049 |
| K17Q | 0.006 | < 0.0001 | 0.003 | < 0.0001 | 1.000 |
| K17A | 0.267 | < 0.0001 | 0.033 | < 0.0001 | 1.000 |
| K17E | 0.022 | 0.189 | 0.007 | < 0.0001 | 0.412 |

[a] *p*-Values for two-way ANOVA with $\alpha$ = .05 and Dunnett's multiple comparison tests (comparisons to degraded Hst5)

**Table S2.** *p*-Values for data evaluating antifungal activity of Hst5 and variants after incubation with and without Saps (Figures 3, 4, 5 and 7).

| Peptide | *p*-Values (Antifungal activity assay)[a] | | | | | |
|---|---|---|---|---|---|---|
| | No Sap (4.7 µg/mL)[b] | Sap1 (4.7 µg/mL) | Sap2 (9.4 µg/mL) | Sap3 (18.8 µg/mL) | Sap9 (18.8 µg/mL) | Saliva (9.4 µg/mL) |
| K13R | 0.0006 | < 0.0001 | 0.9994 | < 0.0001 | 0.0447 | 0.9997 |
| K13L | 0.9996 | < 0.0001 | < 0.0001 | < 0.0001 | < 0.0001 | < 0.0001 |
| K13H | 0.1802 | < 0.0001 | < 0.0001 | < 0.0001 | < 0.0001 | < 0.0001 |
| K13Q | < 0.0001 | < 0.0001 | < 0.0001 | < 0.0001 | < 0.0001 | < 0.0001 |
| K13A | 0.0073 | < 0.0001 | < 0.0001 | < 0.0001 | < 0.0001 | < 0.0001 |
| K13E | < 0.0001 | < 0.0001 | < 0.0001 | < 0.0001 | < 0.0001 | < 0.0001 |
| K17R | 0.486 | < 0.0001 | 0.067 | 0.914 | < 0.0001 | 0.890 |
| K17L | 0.131 | 0.110 | 0.039 | < 0.0001 | < 0.0001 | < 0.0001 |
| K17W | 0.240 | < 0.0001 | 0.039 | 0.002 | < 0.0001 | < 0.0001 |
| K17Q | 0.000 | < 0.0001 | 0.026 | 0.887 | < 0.0001 | < 0.0001 |
| K17A | 0.009 | < 0.0001 | 0.020 | 0.971 | < 0.0001 | < 0.0001 |
| K17E | < 0.0001 | 0.002 | 0.647 | 0.997 | 0.012 | < 0.0001 |

[a] *p*-Values for two-way ANOVA with $\alpha$ = 0.05 and Dunnett's multiple comparison tests (comparisons to degraded Hst5).
[b] Provided concentration is the concentration at which statistical analysis was performed.

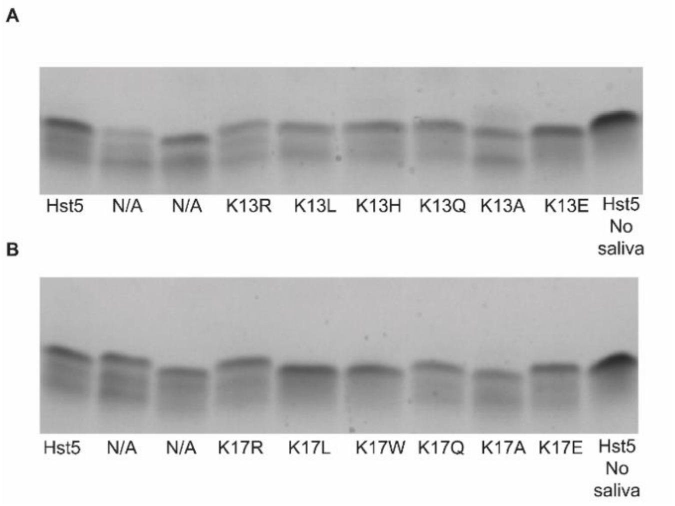

**Figure S5.** Gel images for proteolysis of Hst5 and variants with modification at (A) K13 and (B) K17 by saliva. Irrelevant lanes have been cropped out of the gel images. Lanes marked "N/A" contain variants that are not relevant to this work.

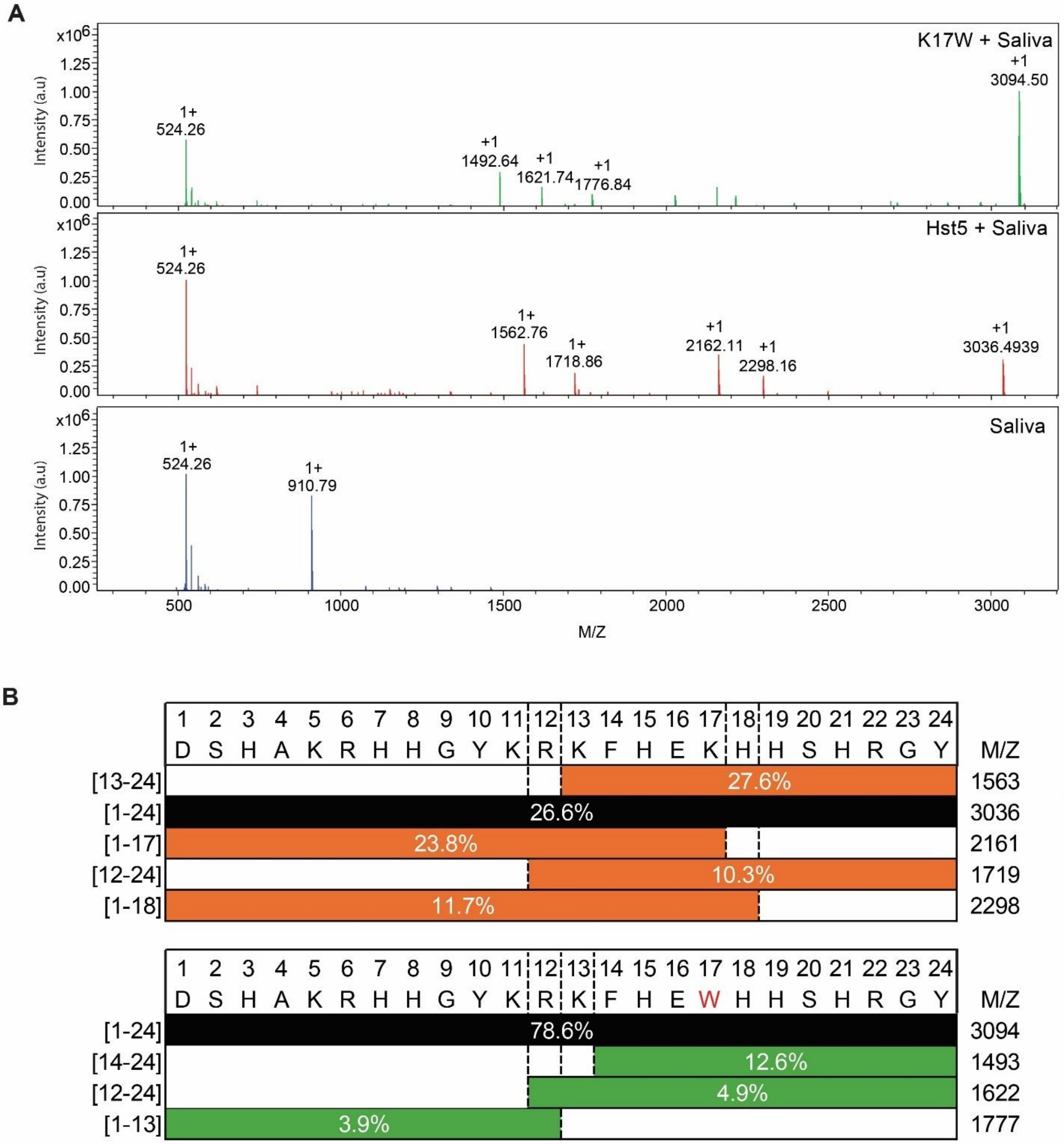

**Figure S6**. Fragments produced by incubation of Hst5 and K17W with saliva. Peptides were incubated with whole saliva for 2 h at 30 °C and analyzed by mass spectrometry. A) Deconvoluted mass spectra for saliva alone, Hst5 incubated with saliva, and K17W incubated with saliva. B) Identified peptide fragments generated by saliva-mediated degradation. Values within each block indicate the relative fraction of that fragment in the sample. Peaks not assigned to peptide fragments represent impurities or signals unrelated to Hst5 or K17W.

**Table S3**. *p*-Values for data evaluating the prevention of biofilm formation by Hst5 and K17W (Figure 8).

| Peptide | *p*-Values (Biofilm prevention assay)[a] | | | |
|---|---|---|---|---|
| | 750 μM | 188 μM | 47 μM | 12 μM |
| K17W | 0.0292 | 0.0004 | 0.2173 | 0.746 |

[a] *p*-Values for two-way ANOVA with $\alpha = 0.05$ and Dunnett's multiple comparison tests (comparison to Hst5).
